# Considerable genetic diversity and structure despite endemism and limited ecological specialization in the Hayden’s ringlet, *Coenonympha haydenii*

**DOI:** 10.1101/2023.05.18.541405

**Authors:** Amy Springer, Zachariah Gompert

**Author notes:** Corresponding author:* Amy Springer, Department of Biology, 5305 Old Main Hill, Utah State University, Logan, UT 84322-5305. **Data Archival Location**: Data and computer code used to analyze these data will be archived in DRYAD (DOI pending). Plant voucher specimens will be pressed, mounted, and archived with the Intermountain Herbarium in Logan, Utah.

## Abstract

Understanding the processes that underlie the development of population genetic structure is central to the study of evolution. Patterns of genetic structure, in turn, can reveal signatures of local adaptation, barriers to gene flow, or even the genesis of speciation. However, it is unclear whether the processes that dominate the development of genetic structure differ in populations with a severely restricted range relative to widespread species. For example, in narrowly endemic species, is population structure likely to be adaptive in nature (e.g., via niche specialization), or rather the result of genetic drift (e.g., isolation by distance)? In this study, we investigated patterns of genetic diversity and structure in the narrow endemic Hayden’s ringlet butterfly. Specifically, we asked to what degree genetic structure in the Hayden’s ringlet can be explained by isolation by distance, barriers to gene flow, and host association. We employed a genotyping-by-sequencing (GBS) approach coupled with host preference assays, Bayesian modeling, and population genomic analyses to answer these questions. Our results suggest that despite their restricted range, levels of genetic diversity in the Hayden’s ringlet are comparable to those seen in non-endemic butterfly species. Hayden’s ringlets showed a strong preference for feeding on grasses vs. sedges, but neither host preference nor potential host availability at sampling sites correlated with genetic structure. We conclude that geography, in the form of barriers to migration and simple isolation by distance, were the major drivers of differentiation in this endemic species.

## Introduction

Genetic structure, or the organization of genetic diversity across geographic space, is a pattern foundational to the study of evolution. It can mark the origin point for speciation (Mayr, 1942; Avise et al., 2000; Harvey et al., 2017), and provides an invaluable tool for elucidating how processes such as genetic drift, gene flow, and natural selection drive evolutionary change (e.g., Wright, 1931; Hey, 2010; Orsini et al., 2013; Lucas et al., 2016; Stankowski et al., 2019; Sendell-Price et al., 2021). In turn, unraveling how evolutionary processes interact and affect the development of genetic structure can provide important insights into the causes and potential consequences of evolution. Analyzing patterns of genetic structure can help reveal migratory routes and patterns of contemporary gene flow (e.g., Gompert et al., 2021; Hemstrom et al., 2022), provide insight into the degree of ecological specialization present across populations (e.g., Nosil et al., 2008; Ferrari et al., 2012; Chaturvedi et al., 2018; Michell et al., 2023), and expose previously unknown patterns of cryptic speciation and hybridization (e.g., Prüfer et al., 2014; Hinojosa et al., 2019). Thus, describing and interpreting patterns of genetic structure remains central to advancing our understanding of evolution.

Patterns of genetic structure are influenced by genetic drift, gene flow, and natural selection (Wright, 1931). But the degree to which the development of genetic structure is dominated by each of these depends heavily on geographic, ecological, and demographic context. When dispersal ability is poor, geographic distance alone can drive population divergence via genetic drift (i.e. isolation by distance, or IBD) (Wright, 1943; Slatkin, 1993). Alternatively, variation in ecological conditions, especially in niche-specialized species, can drive patterns of local adaptation via niche divergence (Funk et al., 2011; Orsini et al., 2013; Driscoe et al., 2019; Luna et al., 2023). Geographic and ecological barriers (e.g., mountains, rivers, host availability) can reduce the homogenizing effects of gene flow on patterns of local adaptation (Irwin et al., 2005; Yuan et al., 2012; Wang and Bradburd, 2014; Mairal et al., 2017; Kopuchian et al., 2020), while demographic factors, including past genetic bottlenecks and source-sink population dynamics, can drive down levels of standing genetic variation and reduce the efficacy of selection vs. drift.

All of these cases—isolation by distance, barriers to gene flow, and local adaptation—can result in population divergence that leads to reproductive barriers and eventually speciation. But which processes play the largest role can be especially important to understand in cases of endemism and conservation concern. Endemism is a condition that would, at first glance, appear to limit the potential for genetic structure to develop. Historically, population genetic theory predicted that endemic species should show low levels of genetic diversity (Frankham, 1997; Soltis and Soltis, 1991). At small population sizes, genetic drift will more readily drive alleles to fixation, leading to loss of diversity over time. This, coupled with a narrowly limited geographic range, would seem to leave little geographic or genetic potential for population structure to arise. But recent studies have shown that this is not necessarily the case (Jiméenez-Mejéıas et al., 2015; Hobbs et al., 2013). Many endemic species, particularly plants, have been found to possess both high levels of genetic diversity (Forrest et al., 2017; Medrano and Herrera, 2008), as well as substantial genetic structure (Turchetto et al., 2016).

The fact that narrow endemic species can show both substantial genetic variation and structure raises questions regarding how patterns of genetic structure develop in these species. Is genetic structure in endemic species likely to be adaptive in nature (see Robitzch et al., 2023), or simply the result of limited gene flow and drift? Do the same processes driving genetic structure in these species also contribute to their range restriction? On one hand, limited population size and low genetic diversity levels might suggest that genetic differentiation in endemic species should be largely driven by isolation by distance and genetic drift. Conversely, evolution of niche specialization can fragment populations and restrict the range a population is able to inhabit. If differentiation is being driven by ecological specialization, then we might predict that local adaptation and niche specialization could also be contributing to the development of endemism. Understanding the nature of population structure in endemic species could help elucidate how range size influences evolutionary processes.

The Hayden’s ringlet, *Coenonympha haydenii*, is a brown Satyrid butterfly found only in mountain meadows and forest clearings of southwestern Montana, southeastern Idaho, and western Wyoming (i.e., the Greater Yellowstone Ecosystem) (Debinski and Pritchard, 2002; Pyle, 1981; Howe and Bauer, 1975; Scott, 1992). A non-migratory narrow endemic, the Hayden’s ringlet is known for both its high local abundances (Caruthers and Debinski, 2006) and weak flying ability (Glassberg, 2001; Kaufman and Brock, 2003). Poor flight could also predispose the Hayden’s ringlet to population divergence via geographic barriers such as mountain ranges. Large local population sizes coupled with poor dispersal ability suggests that enough genetic variation and geographic isolation could exist in this species to result in population structure at small spatial scales. This makes the Hayden’s ringlet an ideal system to investigate processes driving patterns of genetic structure in narrow endemic species.

The Hayden’s ringlet shows potential for ecological divergence. Larvae of *C. haydenii* are thought to feed on one or more species of grasses (family Poaceae) or sedges (family Cyperaceae) (Debinski and Pritchard, 2002; Glassberg, 2001; Feltwell, 1993; Pyle, 1981). It has been suggested that the Hayden’s ringlet may be narrowly distributed simply because they are a remnant species left behind from a larger, pre-glaciation distribution (Pyle, 1981). But the range of the closely-related common ringlet (*Coenonympha tullia*) has expanded dramatically over the past 60 years (Wiernasz, 1983, 1989), while *C. haydenii*’s range, if anything, appears likely to decrease (Debinski et al., 2013; Lotts and Naberhaus, 2021). If the Hayden’s ringlet, like the common ringlet, is a generalist that can feed on a variety of common grasses, why hasn’t the range of the Hayden’s ringlet expanded as well? One possibility is that the Hayden’s ringlet is bounded by the distribution of the host species its larvae feed upon. Female Hayden’s ringlets are thought to be associated more with moist, hydric meadows or bogs rather than dry meadows (Pyle, 1981; Scott, 1992), and species abundance has been shown to decline substantially during periods of drought (Debinski et al., 2013). This is consistent with the idea that *C. haydenii* could be specialized on one or more endemic Yellowstone wetland species like sedges. As such, it is possible natural selection could drive adaptive genetic structure and endemism in this species.

In this study, we characterize patterns of genetic structure in this endemic species and delineate the underlying evolutionary processes that are likely to be driving population structure. Specifically, we asked the following questions: (1) how much genetic diversity and structure exists within the Hayden’s ringlet, (2) to what degree is the development of genetic structure in this endemic species associated with (a) genetic drift and isolation by distance, (b) gene flow and barriers to migration, and/or (c) local adaptation and host availability and preferences.

## Materials and Methods

### Sample Collection

Over the course of two years, we collected adult *C. haydenii* specimens of both sexes from 14 sampling sites across the species’ range (see Fig. 1a). We surveyed for *C. haydenii* presence at two additional locations in the Yellowstone Plateau region (AVP and GLR, see Table S1) and along approximately 10 miles of trail on the John D. Rockefeller Jr. Memorial Parkway, but we only observed a single Hayden’s ringlet across this entire region. Due to low abundance between Yellowstone National Park and Grand Teton National Park, we were unable to collect butterflies from this area. At each of the 14 sites where Hadyen’s ringlet populations were abundant, we collected an average of 27 butterflies per location (see Table 1 for specific sample sizes). Male butterflies were immediately frozen to preserve tissue for subsequent DNA extraction, while females were maintained temporarily in the lab for egg collection and oviposition preference assays and frozen afterwards. Butterfly specimens sampled within Yellowstone and Grand Teton National Parks were collected under permits YELL-2018-SCI-8064, YELL-2019-SCI-8064, GRTE-2018-SCI-0041, and GRTE-2019-SCI-0055.

**Figure 1:**
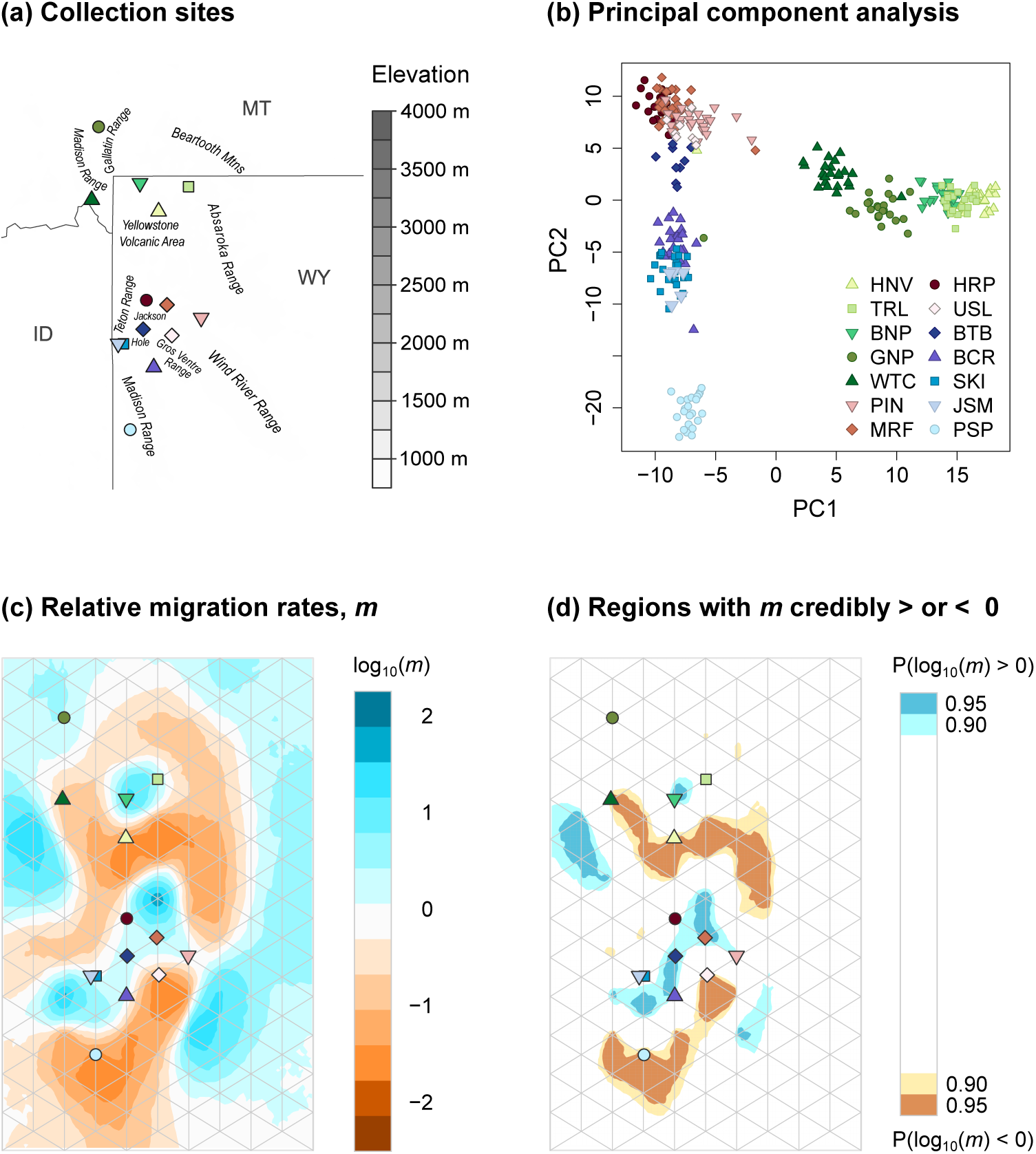
(a) Map of butterfly sampling locations. Each sampling site is depicted as a colored point, the corresponding key for which is shown in panel (b). Elevation contours (in meters) are shown in gray, and major mountain ranges and valley regions within *C. haydenii’s* range are labeled where they occur. (b) Principal component analysis of genotype likelihood estimates from ENTROPY for the 9,313 SNPs. (c) Map of relative migration rates across *C. haydenii’s* range as estimated by EEMS from SNP data. Areas with estimated migration rates lower than expected under isolation by distance (IBD) alone are shown in orange, and those with migration rates higher than expected under IBD are shown in blue. Because EEMS assigns individuals to the nearest vertex on a triangular grid, the locations of populations in the EEMS model do not correspond perfectly to the sampling locations on the geographic map shown in panel (a). (d) Geographic regions with relative migration rates credibly greater or less than zero.

**Table 1:**
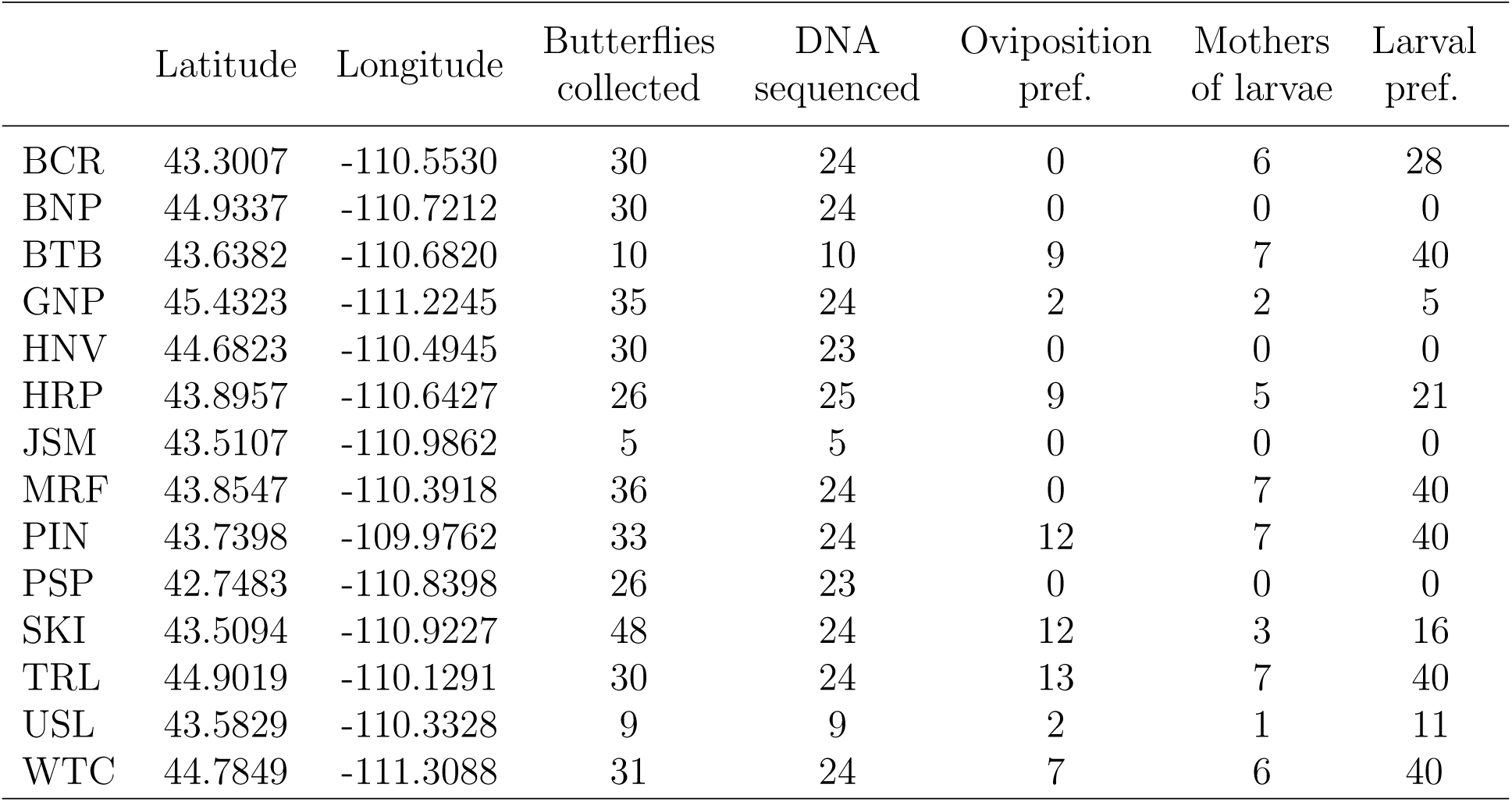
Collection locations and sample sizes for the total number of adult butterflies collected from each site, the total number of specimens from which DNA was extracted and sequenced, the number of female butterflies for which oviposition preference assays were conducted, the number of female butterflies that produced offspring for the larval herbivory assays, and the total number of larvae for which herbivory assays were conducted.

To build a reference collection of potential host species available to *C. haydenii* across their range, we collected plant specimens from 9 of our 14 Hayden’s ringlet butterfly sampling sites. The larvae of the Hayden’s ringlet are suggested to feed generally on grasses (Debinski and Pritchard, 2002; Kaufman and Brock, 2003; Glassberg, 2001), at least some of which may overlap with the host genera used by the closely-related common ringlet butterfly, *C. tullia*. As such, we collected voucher specimens of each unique species of *Poa*, *Stipa*, and *Melica* grasses found in sampling site meadows where Hayden’s ringlets were observed. It has also been suggested that Hayden’s ringlets may be able to feed on sedges (family Cyperaceae) (Feltwell, 1993; Pyle, 1981), so we collected voucher specimens of all species of *Carex* sedges we found as well.

### Host Preference Assays

To determine whether Hayden’s ringlets are more likely to utilize grasses or sedges as a larval host, we conducted both female preference and larval preference assays. For these assays, we chose to compare preference for Hood’s sedge (*Carex hoodii*) vs. Kentucky bluegrass (*Poa pratensis*). Both species are abundant throughout the *C. haydenii’s* range, and represent the two plant families Hayden’s ringlets are hypothesized to feed upon: sedges (Cyperaceae) and grasses (Poaceae). We chose harebell (*Campanula rotundifolia*) as a control group because it is a common herbaceous flower in the area (Craighead, 2005). Harebell is often found growing in meadows in association with grassland communities (Stevens et al., 2012), and thus may realistically be encountered by *C. haydenii* larvae in the wild. Harebell stem leaves are also long and narrow like those of grasses and sedges (Craighead, 2005; McGhan, 2023), which allowed us to control for leaf shape and size during our larval preference assays.

We conducted oviposition preference assays following standard procedures described in Forister et al. (2009). Briefly, adult female butterflies collected from eight of our sampling sites (see Table 1) were placed in plastic cups containing three plant samples each: Hood’s sedge, Kentucky bluegrass, and harebell. Females were maintained in these cups for 72 hours, after which the number of eggs adhered to each species of plant was recorded. Female butterflies were then removed and stored at −80 C until subsequent DNA extraction. After female oviposition preference assays were complete, all eggs laid in the oviposition cups were gently removed from their substrate and stored in vented petri dishes under ambient temperature and light conditions until they hatched (approximately 10 days). Within one day of hatching, we performed larval preference assays following standard protocols (Géomez Jiméenez et al., 2014; Gamberale-Stille et al., 2014; Wang et al., 2017; Gu and Walter, 1999) to assess differences in larval feeding preferences across populations. We tested up to 40 neonate larvae each from 10 of our sampling sites (see Table 1). Larvae were placed in the center of petri dishes equidistant from three 1-cm long leaf segments representing each of our test species (Kentucky bluegrass, Hood’s sedge, and harebell). We took pictures of the leaf tissue flattened between glass slides both before and after the 72 hour herbivory trial with a Canon EOS M6 camera. We then used the program `ImageJ` version 1.52A (Schneider et al., 2012) to trace outlines around each leaf image both before and after herbivory. This allowed us to calculate the amount of surface area lost by each leaf during the herbivory trial. In addition, each leaf was manually assigned a binary value indicating whether signs of herbivory (i.e. jagged leaf margins) were or were not observed in order to better discriminate between area loss that could be attributed to larval herbivory vs. shrinkage due to moisture loss in the leaf over the trial period. See supplement for more information.

### DNA Sequencing, Alignment, and Variant Calling

We used Qiagen DNeasy 96 Blood and Tissue Kits to extract DNA from the thoracic tissue of 287 butterfly specimens representing 14 sampling locations (see Fig. 1 and Table 1). When available, an equal number of male and female specimens were chosen for sequencing from each site. Reduced-representation restriction-fragment based DNA libraries were prepared for genotyping-by-sequencing (GBS) following methods similar to those in Gompert et al. (2014a). Briefly, whole-genome DNA was digested using Mse1 and EcoR1 enzymes, ligated to custom barcode sequences, and amplified via PCR. Barcoded DNA fragments were then pooled across samples, purified, and size-selected using a BluePippin. DNA fragments between 300-450 bp were selected for sequencing. The resulting DNA fragment libraries were sequenced on the University of Texas Illumina HiSeq 4000 sequencing platform. The resulting DNA sequences were first filtered to remove PhiX sequences and poly-G tails. PhiX is a bacterial sequence introduced during HiSeq sequencing as an internal control. We used `SAMtools` version 1.10 and custom scripts to find and remove all reads that aligned to the PhiX reference genome, leaving 347,375,794 individual reads. Barcode sequences were then removed from these remaining reads using custom perl scripts, allowing us to match each DNA sequence to the individual butterfly from which it came.

To date, no reference genome has been published for the Hayden’s ringlet. In the absence of a full reference genome, we constructed a de-novo set of reference contigs for *Coenonympha haydenii* using the program `CD-hit` version 4.8.1 (Li and Godzik, 2006) for aligning our reads. For further details regarding our construction of the reference contig set, see supplement. Reads were aligned to this reference contig set using `BWA` version 0.7.17-r1188 (Li and Durbin, 2009). We used the `BWA aln` algorithm, with the total number of mismatches allowed per read (-n) set to 5, or approximately 0.06% of each read. We set seed length (-l) equal to 20 bp, and the maximum allowed mismatches in the seed sequence (-k) equal to 2.

We identified sites with single nucleotide polymorphisms (SNPs) in our genomic data using `samtools` and `bcftools` version 1.9 (Li et al., 2009). We used the original consensus caller (-c) to call variants, and set the threshold p-value (-p) for accepting variants to 0.01. Variants were then filtered for quality using custom perl scripts. We retained variable sites for which there were at least 2x more reads than the number of individuals we sequenced (i.e., mean coverage per ≥2x), contained a minimum of 10 reads for the alternative allele (to filter out possible sequencing errors), and had a phred-scaled mapping quality >30. We removed variant sites with base quality rank sum test, mapping quality rank sum test, and read position rank sum test p-values less than 0.001, 0.0001, and 0.001 respectively. We also removed any variable sites missing data for 20% or more of the individuals we sequenced. We set a maximum read depth of 8000 to remove possible paralogs/gene families, and removed all sites located less than 2 BP apart along a contig. After quality filtering, we were left with a total of 9,313 SNPs used for downstream analysis.

### Assessing Genetic Diversity and Structure

To measure overall levels of genetic diversity in the Hayden’s ringlet, we calculated both Watterson’s *θ* (*θ_W_*) and nucleotide diversity (*π*). We calculated both diversity statistics using the program `ANGSD` version 0.933-71-g604e1a4 (Korneliussen et al., 2014), which uses the full set of aligned contigs (not our quality-filtered SNP set) to account for uncertainty in the number of segregating sites present. We then calculated per-base pair values of both *θ_W_* and *π* (based on the estimated number of bases sequenced from `ANGSD`) and calculated 95% bootstrap intervals using `R` version 4.2.2.

We used Nei’s *F_ST_*to compare the degree of genetic differentiation across the 14 populations of Hayden’s ringlets we sampled (Nei, 1973). To calculate this, we first used `estpEM` version 0.1 (Soria-Carrasco et al., 2014) to obtain a maximum likelihood estimate of allele frequencies for each SNP (N = 9313) for each population (N = 14) of Hayden’s ringlets we sampled. The program `estpEM` uses an expectation-maximization (EM) algorithm to account for uncertainty in genotypes generated during DNA sequencing (Soria-Carrasco et al., 2014). We set the tolerance level for EM convergence to 0.001, the maximum number of EM iterations to 20, and used our filtered genotype likelihood files split by population as input. With the allele frequency estimates for each population generated by `estpEM`, we then calculated pairwise Nei’s *F_ST_* values for each combination of populations, as well as overall *F_ST_* across all populations. Briefly, we calculated the mean *F_ST_* across all 9313 loci using the formula 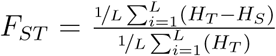 where *H_T_* is the expected heterozygosity at Hardy-weinberg equilibrium for the total population (i.e. across all subpopulations), *H_S_*is the average of the expected heterozygosities within each subpopulation, and L is the number of loci (Lucek et al., 2019). These calculations were completed using custom R code.

Next, we summarized patterns of genetic structure in the Hayden’s ringlet using principal component analysis (PCA) and the program `ENTROPY` version 2.0 (Gompert et al., 2014b; Shastry et al., 2021). `ENTROPY` is a program similar to the admixture model in `STRUCTURE`, but has the added feature of accounting for uncertainty in genotypes as captured by genotype likelihoods. It uses a Bayesian framework to estimate the proportion of a particular individual’s genome that would be derived from each of K hypothetical source populations. The purpose of this in our case was not to estimate the optimal value of K, but rather to assess patterns of gross vs. fine-scale substructure within the species. To this end, we ran `ENTROPY` for all K-values between two and seven using our 9,313-SNP set as input. For each value of K, we ran 10 MCMC chains with a 10,000-step burn-in period, 20,000 sampling iterations, a thinning interval of 5, and a Dirichlet initialization value of 50. We conducted our PCA in `R` using an unscaled genotype correlation matrix produced with genotype estimates generated by `ENTROPY`.

### Causes of Genetic Structure

To identify barriers to migration across *C. haydenii* sites, we used the statistical method Estimating Effective Migration Surfaces (`EEMS`), developed by Petkova et al. (2016). `EEMS` is based on the stepping-stone model of migration, and estimates effective migration rates by comparing the actual degree of genetic differentiation to the expectation under a null isolation-by-distance model. We ran `EEMS` using a grid density of N-demes = 50, 100, and 150 demes. The number of demes corresponds to the number of nodes EEMS produces in the triangular grid to which individual samples can be assigned. For each grid density level, we ran three MCMC chains of 4,000,000 steps each, a burn-in of 2,000,000 steps, and thinning interval of 9999.

To determine the degree to which genetic variation in the Hayden’s ringlet is explained by geographic distance vs. available host plants, we used a Bayesian linear mixed model introduced by Gompert et al. (2014a), which extened a similar maximum likelihood model from Clarke et al. (2002). This model accounts for the lack of independence among sampling site pairs (ie. the genetic distance between populations A vs. B is not independent from the genetic distance between populations A vs. C) (Gompert et al., 2014a). We modeled the effect of host and geographic distance on logit F_ST_ as follows:

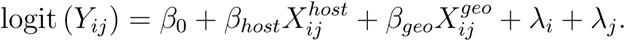

Where *Y_ij_* is mean pairwise F_ST_, 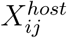 is host dissimilarity (as measured by the Sørensen index), and 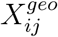 is the geographic distance (calculated as Euclidean distances) between each pair of sites. The Sørensen index measures the number of species shared between two sites as compared to the total number of species present across both sites, with greater weight given to shared than to non-shared species (Hao et al., 2019). The effect of host and geographic distance are represented by *β_host_* and *β_geo_* respectively, and population random effects are represented by *λ_i_* and *λ_j_*. Both geographic and host distance values were centered and standardized prior to running the model to account for differences in unit scale.

To provide additional information about the potential for differences in host use across Hayden’s ringlet populations, we assessed differences in female oviposition preference across populations using a hierarchical Bayesian model. Since female butterflies were given the choice of three plant species for oviposition, we modeled the number of eggs laid on each host multinomially such that P(oviposition on hosts 1:3) ∼ *M* (*P*_(1:3)_*, n*). Each population was allowed its own oviposition preference values, *P*_(1:3)_, in order to test the effect of population on female oviposition preference. The values of *P* from each population were assigned a Dirichlet prior (one for each of the three host plants) with *α* = *π ∗ S*. Here, *π* represents the global preference for each host plant across all populations. Finally, *π* was assigned a Dirichlet hyperprior with *α* = 1, and *S* a uniform hyperprior with lower and upper bounds of 1 and 200, respectively. We fit our model using `rJags` version 4.3.1 (Plummer, 2003, 2013). We ran three MCMC chains of 80,0000 sampling steps each, with a burn-in of 10,000 steps and thinning interval of 50. We checked convergence of the MCMC chains using the Gelman diagnostic (Gelman and Rubin, 1992).

We analyzed larval preference using a complex linear Bayesian model. To determine the total area of each leaf lost during the experimental assay, we calculated the area of the leaf before the herbivory assay (measured in cm^2^) minus the area of the leaf after larvae were allowed to consume leaf tissue for 72 hours. In our model, we assumed leaf area lost during the herbivory assays could be attributed to two main causes: (1) larval feeding, and (2) shrinkage of the leaf tissue due to moisture loss over time. We assumed total leaf area loss to follow a normal distribution with a mean and standard deviation as follows:

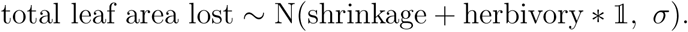

Here, 𝟙 is a binary indicator set equal to 1 if herbivory occurred, and equal to 0 if no herbivory occurred. Thus, in cases where herbivory occurred, mean leaf area lost was defined as the sum of shrinkage plus larval herbivory. If no herbivory occurred, mean leaf area lost was defined as shrinkage only. We defined herbivory as following a normal distribution where the mean and standard deviation were allowed to vary by each unique plant host species x butterfly population combination. Shrinkage was defined as following a normal distribution where the mean and standard deviation were allowed to vary by host plant only since the population each caterpillar was obtained from should have no effect on the amount of moisture lost by each leaf over time during the assay. We wrote this model in the language STAN (Stan Development Team, 2022b) and implemented it using the R-interface RStan version 2.21.5 (Stan Development Team, 2022a). We used a warm-up period of 15,000 steps and ran the model for 30,000 MCMC steps.

## Results

### *C. haydenii* shows strong preference for grass host

We saw a strong preference for oviposition on Kentucky bluegrass across all populations of *C. haydenii* we assessed (see Fig. 4). If female butterflies distributed their eggs randomly across available substrates, then we would expect each of the three host plants to receive 33% of all eggs laid, on average. In contrast, the median global preference for oviposition on Kentucky bluegrass was 57% (95% CI 47-67%; posterior probability [p.p.] percent oviposition on *P. pratensis >* 33% > 0.99). Preference for oviposition on Kentucky bluegrass was credibly higher than expected by chance for all eight populations we assayed (p.p. percent oviposition on *P. pratensis >* 33% > 0.98 for all populations). The global preference for oviposition on Hood’s sedge, *Carex hoodii*, was credibly lower than expected by chance (p.p. percent oviposition on *C. hoodii <* 33% = 0.96, median oviposition 24%, 95% CI 17-35%). Median global preference for oviposition on harebell, our control species, was the lowest at only 17% (95% CI 10-26%; p.p. preference for *C. rotundifolia <* 0.33 > 0.99), 16 percentage points lower than expected if butterflies distributed their eggs randomly across available substrates.

While all populations of Hayden’s ringlets we sampled showed a preference for oviposition on Kentucky bluegrass, the degree of preference for oviposition on both grass and sedge varied across populations. Female butterflies from BTB showed the strongest overall preference for Kentucky bluegrass, laying 74% of their eggs on *P. pratensis* (95% CI 61-87%). Meanwhile, WTC and HRP showed the lowest rates of oviposition on *P. pratensis*, laying only 51% of their eggs on the grass host (95% CI 37-63% and 33-65% respectively). Preference for oviposition on *Carex hoodii* also varied, with PIN showing the highest rates of oviposition on sedge (33.1%, 95% CI 22-46%), and TRL showing the lowest at 17% (95% CI 10-25%). Comparing overall preference across populations, we found that both the TRL and BTB populations showed credibly higher rates of oviposition on *Poa pratensis* than the PIN, HRP, and WTC populations (p.p. > 0.99 for all six comparisons), and PIN showed credibly higher percent oviposition on *Carex hoodii* than BTB (95% CI for PIN – BTB > 0; % oviposition on carex for BTB = 8-27%, while % oviposition on carex for PIN = 22-46%). We found that three populations (BTB, TRL, and WTC) showed a credibly greater preference for oviposition on Kentucky bluegrass vs. Hood’s sedge (95% CI for Pref KB – Pref CH > 0 for all three populations). While all populations but two showed a credible difference in rates of oviposition on KB vs. harebell (GNP and HRP were the exceptions), only one population showed a credible difference in rates of ovipositon on carex vs. harebell (PIN, difference = 18 percentage points, 95% CI = 1-36%).

As with oviposition, Hadyen’s ringlet larvae showed a strong preference for Kentucky bluegrass, *Poa pratensis*. The species-level preference for *Poa pratensis* produced by the Bayesian model was 71% (95% CI 64%-79%), meaning we would expect a randomly sampled group of Hayden’s ringlet larvae to consume a mean of 71% *Poa pratensis* grass when offered a choice of *Poa pratensis, Carex hoodii*, and emphCampanula rotundifolia. Unlike in the female oviposition assays, no harebell herbivory was observed from any of the larvae we assayed. All populations we assayed showed a trend toward consuming more grass (*Poa pratensis*) than sedge (*Carex hoodii*), with every population consuming credibly more grass than sedge (P(consumed more grass than sedge) > 0.99) except SKI (P(consumed more grass than sedge) = 0.85).

The total amount of leaf tissue consumed by larvae varied considerably across populations, with some populations consuming more leaf tissue overall than others (see Fig. 4b). Total leaf tissue consumption by population ranged from a minimum of 3.2 mm^2^ (HRP) to a maximum of 4.6 mm^2^ (SKI), with 25 out of 45 population pairs showing credible differences in total larval feeding. Due to these differences in overall larval feeding levels, we assessed differences in host preference across populations as differences in the proportion of grass vs. sedge leaf tissue consumed. For these calculations, we set any negative values in the MCMC chains to zero (as negative herbivory is not a realistic assumption) and removed any NAs that resulted during the proportion calculations. Because no herbivory was observed on the control host, *C. rotundifolia*, we calculated the proportion of grass (or sedge) consumed as simply the total grass (or sedge) tissue consumed, in mm^2^, divided by the sum of grass plus sedge tissue consumed.

The proportion of each host species eaten varied considerably by population. BCR and HRP showed the greatest preference for *Poa pratensis*, consuming 100% grass (95% CI 87-100% and 75-100% respectively) and 0% *Carex hoodii* sedge (95% CI 0-13% and 0-25% respectively). SKI, meanwhile, showed the lowest degree of preference, consuming 56.4% *Poa pratensis* grass (95% CI 44-71%) vs. 44% *Carex hoodii* sedge (95% CI 29-56%). We saw credible differences in herbivory preference for 21 out of 45 pairwise population comparisons. The pairs with the greatest differences in preference were BCR vs. SKI and HRP vs. TRL, with BCR and HRP consuming 42.1 (95% CI 27-55) and 41.9 (95% CI 17-55) percentage points more *Poa pratensis* grass and 42.1 (95% CI 27-55) and 41.9 (95% CI 17-55) percentage points less *Carex hoodii* than SKI and TRL respectively.

### Moderate genetic diversity and population structure exist in the endemic Hayden’s Ringlet

Estimates of *θ_W_*across populations of the Hayden’s ringlet varied from a low of 0.00280 (95% bootstrap interval 0.00277-0.00282) at JSM to a high of 0.00360 (95% bootstrap interval 0.00359-0.00362) at BNP (see Table 2). Estimates of genetic diversity (*θ_π_*) were similar, ranging from 0.00284 at JSM (95% bootstrap interval 0.00281-0.00288) to 0.00344 at USL (95% bootstrap interval 0.00342-0.00347) (see Table 2). Genetic structure across sites was moderate but notable, with an overall *F_ST_* of 0.10. Pairwise *F_ST_* values (see Table 3) ranged from 0.0181 to 0.1191. The population pairs that showed the highest degree of genetic differentiation were USL vs. JSM (*F_ST_*= 0.1191) and USL vs. PSP (*F_ST_*= 0.1071). Meanwhile, the least-differentiated population pairs were TRL vs. BNP (*F_ST_* = 0.0181) and HRP vs. MRF (*F_ST_* = 0.0186). JSM and SKI, which are located very closely in geographic space (∼5 km apart, see Fig. 1a) nevertheless showed a degree of differentiation comparable to population pairs much further apart in geographic space (*F_ST_* = 0.0609). Principal component analysis shows individuals clustering clearly by sampling site (see Fig. 1b), indicating that enough genetic structure exists within this species to identify roughly where in geographic space a particular butterfly originated from based on the DNA sequence data we generated. In particular, we saw that principal component 1 separates the northern Hayden’s ringlet populations from southern populations, while principal component 2 separates the southern populations of Hayden’s ringlets along a NE to SW gradient. The PCA does not perfectly mirror our sampling locations, but nevertheless indicates isolation by distance.

**Table 2:** Watterson’s *θ* (*θ_W_*) and nucleotide diversity (*π*) with 95% bootstrap confidence intervals.

| Population | $\theta_w$ | $\pi$ |
| --- | --- | --- |
| BCR | 0.00348 (0.00347, 0.00349) | 0.00333 (0.00331, 0.00334) |
| BNP | 0.00360 (0.00359, 0.00362) | 0.00332 (0.00329, 0.00334) |
| BTB | 0.00311 (0.00310, 0.00313) | 0.00312 (0.00309, 0.00314) |
| GNP | 0.00347 (0.00345, 0.00348) | 0.00327 (0.00325, 0.00329) |
| HNV | 0.00301 (0.00299, 0.00302) | 0.00298 (0.00296, 0.00300) |
| HRP | 0.00335 (0.00334, 0.00337) | 0.00323 (0.00321, 0.00325) |
| JSM | 0.00280 (0.00277, 0.00283) | 0.00284 (0.00281, 0.00288) |
| MRF | 0.00353 (0.00351, 0.00354) | 0.00338 (0.00336, 0.00340) |
| PIN | 0.00330 (0.00329, 0.00331) | 0.00327 (0.00325, 0.00329) |
| PSP | 0.00327 (0.00326, 0.00329) | 0.00319 (0.00317, 0.00321) |
| SKI | 0.00339 (0.00338, 0.00341) | 0.00313 (0.00311, 0.00315) |
| TRL | 0.00349 (0.00348, 0.00350) | 0.00321 (0.00319, 0.00322) |
| USL | 0.00324 (0.00322, 0.00326) | 0.00344 (0.00342, 0.00347) |
| WTC | 0.00330 (0.00329, 0.00331) | 0.00308 (0.00306, 0.00310) |

**Table 3:**
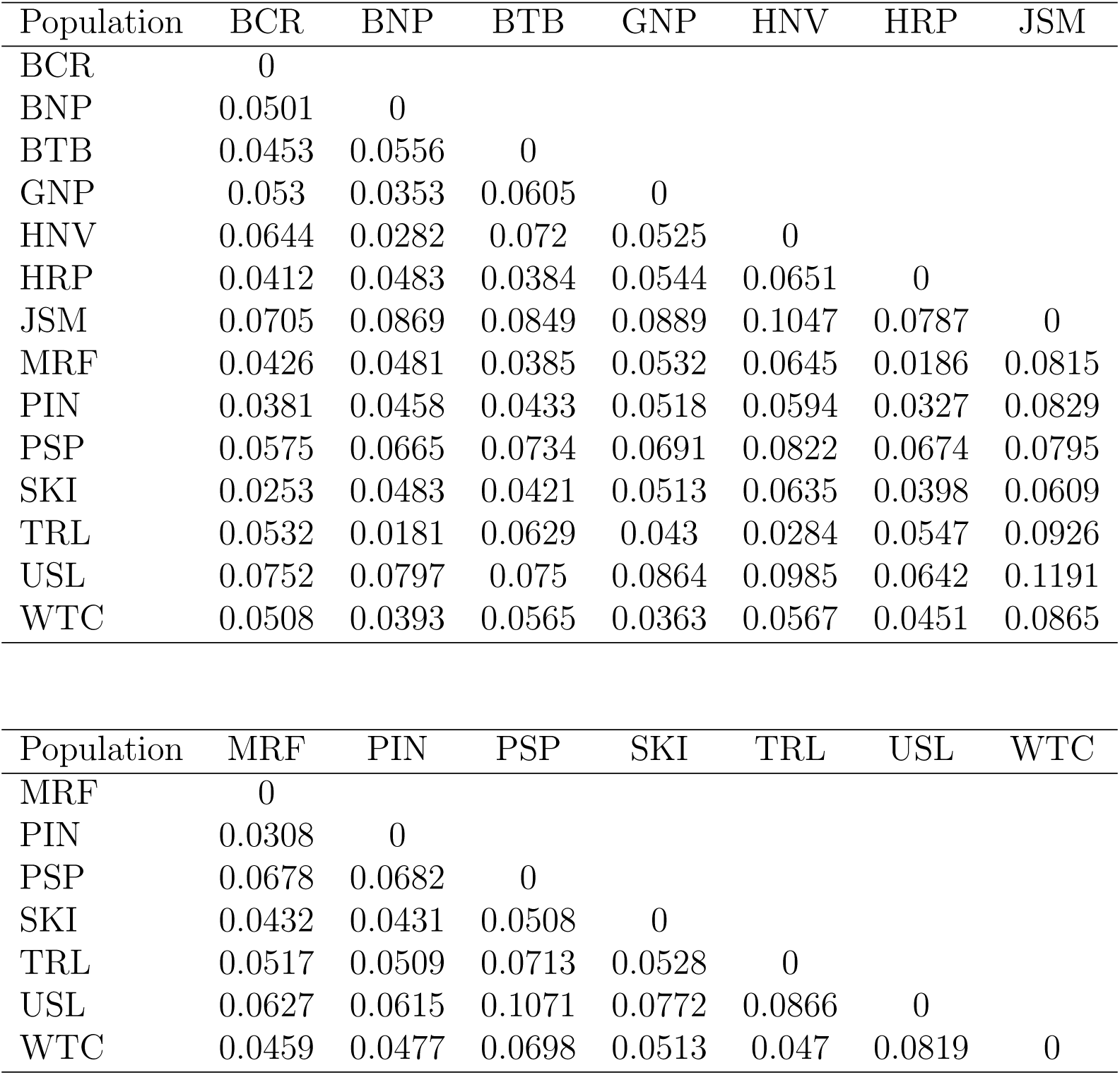
Pairwise F_ST_ values calculated from EEMS genotype estimates.

| Population | BCR | BNP | BTB | GNP | HNV | HRP | JSM |
| --- | --- | --- | --- | --- | --- | --- | --- |
| BCR | 0 |  |  |  |  |  |  |
| BNP | 0.0501 | 0 |  |  |  |  |  |
| BTB | 0.0453 | 0.0556 | 0 |  |  |  |  |
| GNP | 0.053 | 0.0353 | 0.0605 | 0 |  |  |  |
| HNV | 0.0644 | 0.0282 | 0.072 | 0.0525 | 0 |  |  |
| HRP | 0.0412 | 0.0483 | 0.0384 | 0.0544 | 0.0651 | 0 |  |
| JSM | 0.0705 | 0.0869 | 0.0849 | 0.0889 | 0.1047 | 0.0787 | 0 |
| MRF | 0.0426 | 0.0481 | 0.0385 | 0.0532 | 0.0645 | 0.0186 | 0.0815 |
| PIN | 0.0381 | 0.0458 | 0.0433 | 0.0518 | 0.0594 | 0.0327 | 0.0829 |
| PSP | 0.0575 | 0.0665 | 0.0734 | 0.0691 | 0.0822 | 0.0674 | 0.0795 |
| SKI | 0.0253 | 0.0483 | 0.0421 | 0.0513 | 0.0635 | 0.0398 | 0.0609 |
| TRL | 0.0532 | 0.0181 | 0.0629 | 0.043 | 0.0284 | 0.0547 | 0.0926 |
| USL | 0.0752 | 0.0797 | 0.075 | 0.0864 | 0.0985 | 0.0642 | 0.1191 |
| WTC | 0.0508 | 0.0393 | 0.0565 | 0.0363 | 0.0567 | 0.0451 | 0.0865 |

Table 3: (*continued*)
| Population | MRF | PIN | PSP | SKI | TRL | USL | WTC |
| --- | --- | --- | --- | --- | --- | --- | --- |
| MRF | 0 |  |  |  |  |  |  |
| PIN | 0.0308 | 0 |  |  |  |  |  |
| PSP | 0.0678 | 0.0682 | 0 |  |  |  |  |
| SKI | 0.0432 | 0.0431 | 0.0508 | 0 |  |  |  |
| TRL | 0.0517 | 0.0509 | 0.0713 | 0.0528 | 0 |  |  |
| USL | 0.0627 | 0.0615 | 0.1071 | 0.0772 | 0.0866 | 0 |  |
| WTC | 0.0459 | 0.0477 | 0.0698 | 0.0513 | 0.047 | 0.0819 | 0 |

Admixture analysis (Fig. 2) showed the presence of meaningful structure across populations of Hayden’s ringlets across multiple levels of K. The most prominent pattern was a clinal split between the northern and southern populations of Hayden’s ringlets at K=2. Higher values of K revealed additional substructure within the species. At K = 3, `ENTROPY` split the southern populations of Hayden’s ringlets along a North-South axis. In particular, we saw the southernmost population of Hayden’s ringlets, PSP, being separated from the remainder of the populations. Similarly, K = 4 split the northern populations across a roughly West-East axis, separating northern populations east of the Gallatin mountain range (BNP, HNV, TRL) from those west of this range (GNP, WTC). Higher levels of K continued to refine the northeast-to-southwest clinal pattern seen across the southern populations of Hayden’s ringlets. A small number of individual butterflies (specifically from BCR, MRF, GNP, and HNV) showed ancestry values that differed considerably from both the typical values of their own population, as well as those of other populations we surveyed. This suggests that these individuals could be migrants or of mixed ancestry. Overall, our admixture analysis suggests that the greatest degree of genetic differentiation in the Hayden’s ringlet exists between northern and southern populations, with additional substructure occurring within those locations.

**Figure 2:**
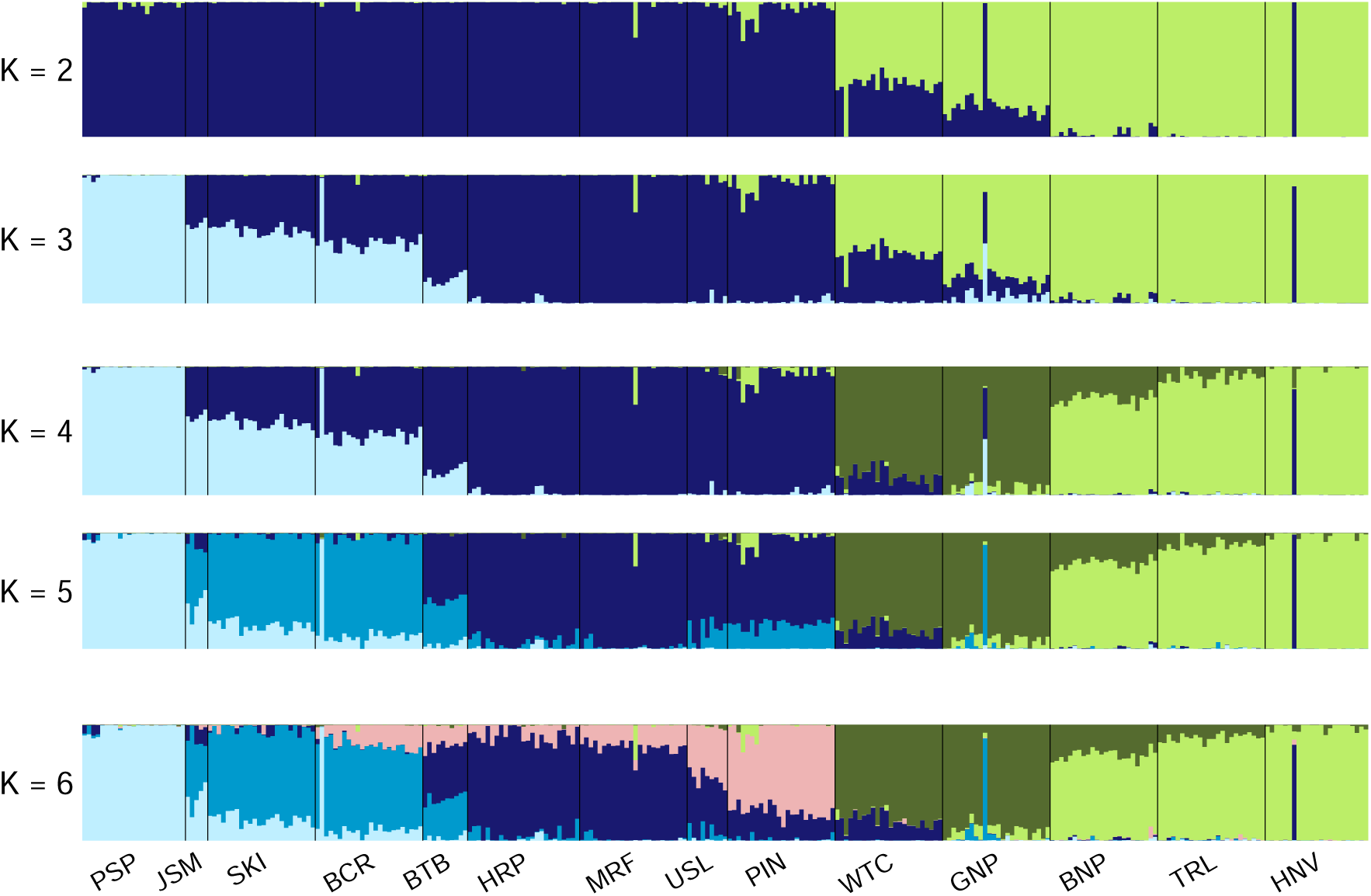
Estimated admixture proportions assuming individuals were sampled from K = 2 through K = 6 hypothetical source populations. Each vertical segment on the barplot represents the estimated ancestry of an individual butterfly, with the proportion of each color in the segment representing the proportion of that butterfly’s genome estimated to have been inherited from each of the K putative source populations. Individuals are grouped along the x-axis by population, with populations demarcated by vertical black bars.

### Isolation by distance and barriers to migration, but not host preference, contribute to population structure in *C. haydenii*

We saw a strong signal for isolation by distance (See Figure 3) in the Hayden’s ringlet, with *β_geo_* = 0.31 (95% CI = 0.24-0.38). The Mantel test also showed a strong relationship between geographic and genetic distances in the Hayden’s ringlet, with a Mantel correlation value of R = 0.7. In addition to simple isolation by distance (IBD), `EEMS` analysis showed several geographic areas with credibly increased or reduced relative migration rates (see Fig. 1c and 1d). Results for each of the three chains for grid sizes of 50, 100, and 150 were similar (see Fig. S1). There were several geographic areas within *C. haydenii*’s range where genetic differentiation among populations was either lower or higher than expected under an IBD model alone. This suggests that certain regions of the physical landscape are easier or more difficult for Hayden’s ringlets to migrate across. In particular, we saw a region of credibly reduced relative migration rates separating the northern and southern populations in our study, consistent with results from PC1 of the principal component analysis (see figure 1d and 1b). This geographic region of credibly reduced gene flow produced by the `EEMS` model corresponds to the location of the Yellowstone plateau, roughly following the southern edge of the geothermally active Yellowstone volcanic area (see Fig. 1a). There was also a region of credibly increased relative migration connecting the majority of the southern populations of Hayden’s ringlets except PSP, the southernmost population. This region of increased connectivity among southern Hayden’s ringlet populations follows the river valley region known as Jackson hole, a low-elevation region between the Teton and Gros Ventre mountain regions (see Fig. 1a). The southernmost population (PSP), which showed credibly lower levels of gene flow with the remaining ringlet populations than expected under a simple IBD model, is separated from the Jackson hole valley region by the Wyoming mountain range.

**Figure 3:**
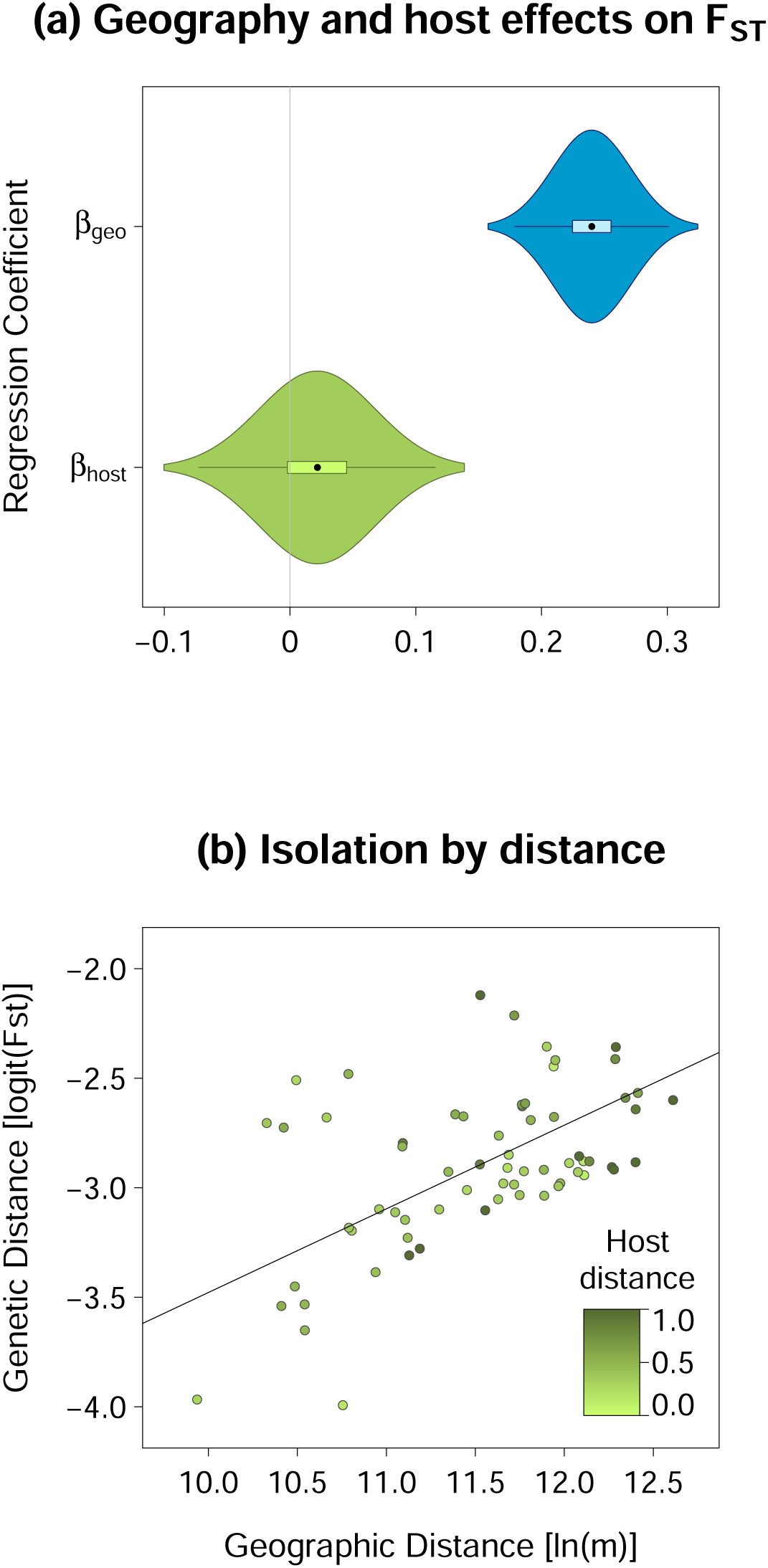
Posterior distributions for the regression coefficients in the Bayesian model estimating the degree to which host distance and geographic distance predict genetic distance (logit F_ST_). (b) shows the linear relationship between genetic distance (as logit F_ST_) vs. geographic distance (ln[meters]) for each pairwise combination of source populations except X and Z.

**Figure 4:**
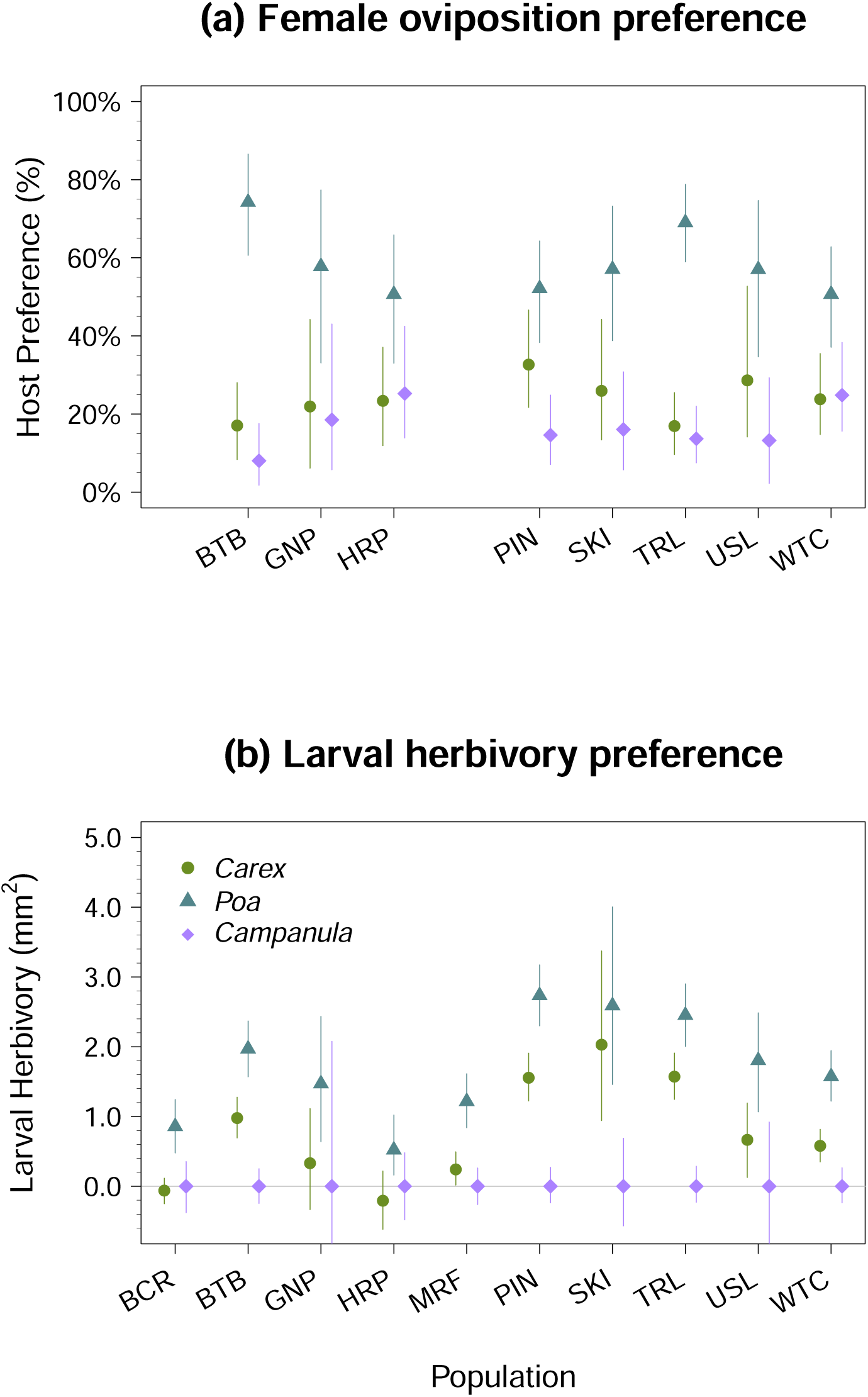
(a) Oviposition preference for female *C. haydenii* from 8 of our sampling sites. (b) Differences in larval herbivory across hosts for each population assayed. The expected total leaf tissue consumption for a caterpillar from a given population is shown on the y-axis. Leaves offered to larvae during the hervivory assays had a mean surface area of 15.7 mm^2^.

Finally, we saw no measurable effect of host distance (as measured by the Sørensen index) on degree of genetic differentiation in the Hayden’s ringlet (see Fig. 3). The credible interval for *β_host_*overlapped zero (P(*β_host_ >* 0) = 73%), indicating there was no credible effect of the presence of Poa, Stipa, Melica, and Carex species on genetic differentiation in the Hayden’s ringlet.

## Discussion

In this study, we assessed patterns of genetic diversity and structure in the endemic Hayden’s ringlet. We also assessed patterns of oviposition and larval host preference, and used Bayesian methods and EEMS modeling to assess the role of isolation by distance, barriers to gene flow, and host availability and preference contribute to population structure in this species. Our results indicate that despite range restriction, the Hayden’s ringlet shows genetic diversity levels comparable to other more widely-distributed species. The Hayden’s ringlet also appears to uniformly prefer grass (*Poa pratensis*) over sedge (*Carex hoodii*), but this host association is unlikely to be driving endemism or patterns of population structure. Instead, we found that both isolation by distance and barriers to gene flow were most closely correlated with genetic distances in this species. We discuss the implications of these results in more detail below.

### Endemism not associated with notable genetic diversity reduction in the Hayden’s ringlet

Despite its restricted distribution, the Hayden’s ringlet showed levels of genetic diversity comparable to other similar butterfly species. The average genetic diversity across Hayden’s ringlet populations we sampled was *θ_π_* = 0.003. Studies on other non-migratory butterfly species with poor dispersal (*Leptidea* sp.; *Lycaeides melissa*; *Parnassius mnemosyne*) reported similar values, with genetic diversity levels between *θ_π_* = 0.001-0.005 (Talla et al., 2019; Gompert et al., 2014b; Talla et al., 2023). Notably, each of the aforementioned species is widely distributed. This suggests that while low, genetic diversity levels in the Hayden’s ringlet are still well within the range of expectation for comparable, non-endemic species. In contrast, both migratory monarchs (*Danaeus plexippus*) and non-migratory *Heliconius* sp. showed comparatively high genetic diversity (*θ_π_* = 0.01-0.06 and 0.020-0.28, respectively (Talla et al., 2020; Hemstrom et al., 2022; Martin et al., 2016; Kryvokhyzha, 2014). Migratory butterfly species have been shown to harbor higher levels of genetic diversity than non-migratory species in general, possibly due to greater population sizes and connectivity (Garćıa-Berro et al., 2023), so the substantial difference in genetic diversity between monarchs and Hayden’s ringlets is not unexpected. However, *Heliconius* species are both non-migratory and have low dispersal ability (Kronforst and Fleming, 2001), so why this species group shows far higher genetic diversity levels than reported in other non-migratory species is unclear.

Many butterfly species have wide distributions, but are locally rare. The Hayden’s ringlet, by contrast, is narrowly restricted in range, but locally prolific. Within their range, Hayden’s ringlets are often so abundant they are the most common butterfly species surveyed (Caruthers and Debinski, 2006). High local abundances in the Hayden’s ringlet could be one factor contributing to the maintenance of genetic diversity in this species. Conversely, poor dispersal (as seen in *Lycaeides melissa* and *Parnassius mnemosyne*) (Gompert et al., 2010; Talla et al., 2019; Gorbach and Kabanen, 2010)—and consequently higher levels of genetic drift within isolated populations—could be reducing population-level nucleotide diversity estimates in more widely-distributed species, especially if local abundances are low. In particular, the widely-distributed *Lycaeides melissa* is known for patchy distributions and metapopulation dynamics (Scott 1992; Gompert et al. 2010; but also see Guiney et al. 2010), which may contribute to lower population-level diversity despite their wide overall range. In all, the similarity in diversity levels between the Hayden’s ringlet vs. widely-distributed species suggest this is yet another case where narrow endemism is not associated with a reduction in genetic diversity. This adds to a growing body of research showing that even highly endemic species can still harbor substantial genetic diversity (Forrest et al., 2017; Medrano and Herrera, 2008; Robitzch et al., 2023; Hobbs et al., 2013; Jiméenez-Mejéıas et al., 2015). In the case of non-migratory butterflies, it appears that local abundances and dispersal capacity may be just as—if not more—critical in determining the amount of genetic diversity maintained than endemism.

### Geography informs patterns of population genetic structure in the Hayden’s ringlet

We saw clear evidence of population structure across the range of the Hayden’s ringlet. The strongest signal of genetic differentiation was a geographic split between northern and southern populations of *C. haydenii*, with additional genetic substructure occurring within each of these groups. We also saw a clear pattern of isolation by distance, with genetic divergence (logit(F_ST_)) increasing by 0.31 standard deviations for every 1 standard deviation increase in geographic distance (m). The correlation coefficient between geographic and genetic distance from the Mantel test was R = 0.7, substantially higher than correlation values seen in other non-migratory species. By comparison, Mantel test correlations for the Langue’s metalmark (*Apodemia mormo langei*), heath fritillaries (*Melitaea athalia* and *Melitaea celadussa*), and checkerspots (*Euphydryas aurinia* and *Euphydryas editha*) ranged between R = 0.39 and R= 0.53 (Dupuis et al., 2018; Tahami et al., 2021; Mikheyev et al., 2013). This suggests that isolation by distance is able to explain a greater degree of the population structure observed in the Hayden’s ringlet than in other similar species. If the Hayden’s ringlet is indeed a generalist feeder, it may show less local adaptation than one might find in a butterfly species specialized on just one or a few hosts. The high correlation between genetic and geographic distances in the Hayden’s ringlet suggests much of the population structure observed can be attributed not to ecological divergence, but to genetic drift and limited dispersal.

Despite the clear patterns of genetic structure present in this species, F_ST_ values between populations of Hayden’s ringlets were low to moderate. The scale of differentiation we observed is consistent with fine-to moderate-scale genetic population structure (F_ST_ between 0.01-0.2) seen in other non-migratory butterfly species (Talla et al., 2019; Pertoldi et al., 2021; Talla et al., 2023; Hinojosa et al., 2023), and on average greater than in migratory species like monarchs (F_ST_ = 0.0001) (Talla et al., 2020). While the F_ST_ values we observed may be considered low in other species, in many cases F_ST_ values between species of butterflies are not considerably greater than what we found within populations of the Hayden’s ringlet (i.e. Talla et al., 2019; Tahami et al., 2021), and in some cases variation within butterfly species is higher than that observed between species. For example, in the El Segundo blue (*Euphilotes battoides allyni*), F_ST_ among populations of the same species ranged from 0.1 to 0.5 (Dupuis et al., 2020), while in heath fritillaries, F_ST_ between two nominal species (*Melitaea celadussa* and *Melitaea athalia*) was only 0.1-0.2 (Tahami et al., 2021). F_ST_ values are affected by both genetic diversity levels (the lower the diversity, the higher F_ST_,) and by sample size (the lower the sample size, the higher the F_ST_), our results are clearly in-line with results from other butterfly species, and consistent with expectations for a non-migratory species with limited dispersal ability.

We saw several geographic regions with credibly increased or reduced relative migration rates in the Hayden’s ringlet. The largest of these was a wide region of credibly reduced relative gene flow between northern and southern *C. haydenii* populations corresponding to the southern border of the Yellowstone plateau and John D. Rockefeller, Jr. Memorial Parkway. Despite having visited two additional sites (Avalanche Peak AVP, Grassy Lake Reservoir GLR) and surveyed approximately 10 miles of trail in this region (see Table S1), we found no viable populations of Hayden’s ringlets connecting our northern and southern sampling sites. Much of the habitat in this region consisted of dense lodgepole pine monocultures growing in previous burn sites (Parmenter et al., 2003; Turner and Simard, 2017; Rothermel, 1994). Hayden’s ringlets prefer open grassy meadows and sunny forest edges (Debinski and Pritchard, 2002; Kaufman and Brock, 2003), so this densely-forested region could present an ecological barrier to migration. Regardless, the fact that our field observations are consistent with the results from our EEMS model suggests that this geographic region presents a true barrier to gene flow for the Hayden’s ringlet. Interestingly, the geographic split we found between northern and southern *C. haydenii* populations corresponds to a similar boundary observed between northern *Lycaeides idas* populations and southern, admixed *Lycaeides* (Gompert et al., 2010, 2012). This suggests that a combination of geographic (elevation; mountain ranges) and ecological (forest type) conditions present in the John D. Rockefeller, Jr. Memorial Parkway may present a barrier to gene flow more generally, and could apply to other non-migratory butterfly species in the greater Yellowstone ecosystem as well.

Despite being a non-migratory species known for poor flight (Kaufman and Brock, 2003; Glassberg, 2001), we nevertheless saw evidence of long-distance dispersal in *C. haydenii*. Several individuals in our admixture analysis matched neither the population from which they were sampled, nor any other population we sampled. In particular, one individual each from MRF, GNP, and HNV in our admixture plots did not match the admixture proportions of any other butterflies we sampled. These individuals appear to be either of hybrid origin or migrants from an area we did not sample. One individual from BCR, on the other hand, appears to be a migrant from PSP (or near PSP). The distance between PSP and BCR is over 65 km, indicating that long-distance dispersal does occur in *C. haydenii* at least occasionally. Hayden’s ringlets are notoriously poor fliers (Glassberg, 2001; Kaufman and Brock, 2003), so we expect typical dispersal distances in the Hayden’s ringlet to be similar to those reported for other poor dispersers like *Lycaeides melissa*, *Parnassius* sp., and *Heliconius* sp. (Gompert et al., 2010; Gorbach and Kabanen, 2010; Kronforst and Fleming, 2001), which rarely disperse further than 2 km during their lifetime. We suggest that the instances of long-distance dispersal we report here are likely a result of rare gene flow events such as butterflies being blown long distances during adverse weather conditions. But as even small amounts of gene flow are sufficient to erase patterns of genetic differentiation, these occasional long-distance dispersal events likely still play a role in determining the magnitude of population genetic structure present in this species.

### Strong preference for grass host, but no evidence of ecological divergence across populations in the Hayden’s ringlet

We observed strong oviposition and larval herbivory preference for Kentucky bluegrass (*Poa pratensis* over Hood’s sedge *Carex hoodii* in *C. haydenii*. Preference for grass was both strong and remarkably consistent, with all populations showing a credible preference for *Poa* in both oviposition and herbivory assays with the exception of SKI. While it has been previously suggested that Hayden’s ringlets might feed on sedges due to their association with bogs and hydric habitats (Pyle, 1981; Scott, 1992), our evidence overwhelmingly points to grasses as being the preferred host of the Hayden’s ringlet. However, the fact that larvae did often feed on both the sedge and grass host, while completely refusing the control host, suggests that Hayden’s ringlets are not likely to be narrowly host-specialized, but generalized feeders like their congener the common ringlet, *C. tullia* (Scott, 1992; Debinski and Pritchard, 2002). This is consistent with preliminary host acceptance data we collected which showed that Hayden’s ringlet larvae will consume tissue from many genera of grasses and sedges including *Stipa*, *Carex*, *Poa*, *Phleum*, and *Elymus*. Anecdotal evidence also suggests that Hayden’s ringlet larvae can be reared to adulthood on *Carex* species (Stout, 2017)(personal communication, Todd Stout, 2017, HOW TO CITE?), which would indeed suggest that the Hayden’s ringlet is a broad generalist given their strong preference for *Poa*. That said, our study only compared only a single species of sedge with a single species of grass. It is possible these species alone are not sufficient to provide a full picture of *C. haydenii’s* preference for grasses vs. sedges. Additional work is needed to further elucidate the degree of host specificity and preference in *C. haydenii*.

While the degree of preference for *Poa* varied credibly across populations, we saw no evidence of host-associated genetic differentiation across populations in the Hayden’s ringlet. Neither sampling site host species composition nor differences in larval preference were predictive of genetic distances among Hayden’s ringlet populations in our study. If the Hayden’s ringlet is in fact a generalist feeder, and host use does not substantially impact larval fitness, then the composition and abundance of potential host species may have a limited effect on genetic differentiation. This could explain the absence of host-associated population structure we observed in this species. But how then do we interpret the phenotypic variation in host preference among populations we observed? It is possible the variation we saw reflects true variation for preference that exists among Hayden’s ringlet populations in the wild. It is also possible that because of the confounding of geographic and ecological distance, the strong signal for isolation by distance we observed is obscuring any signal of ecological differentiation. However, laboratory experiments must always be interpreted with caution with regard to their applicability in the field. In this case, we note that the Hayden’s ringlet populations that showed the highest degree of herbivory preference also happened to be the populations that consumed the least total amount of plant material. Because our preference measure was scaled by total tissue consumed, the lower the total level of consumption, the more sensitive (and stochastic) our preference measure will be to small differences in herbivory. In other words, when total consumption is low, each bite of tissue a larva consumes will have a proportionally larger impact on preference than that same bite of tissue in a case where total consumption is high. Thus, in cases where total herbivory was low, herbivory preferences have the potential to appear exaggerated compared to cases where larvae ate a greater amount of total leaf tissue.

Given the absence of a signal for either strong host specialization or host-associated genetic differentiation, we are left with one major question: what is driving endemism in the Hayden’s ringlet? Since we only assayed two species of potential hosts, we cannot definitively say that host-specialization is not a driver of genetic differentiation and endemism in the Hayden’s ringlet. But preliminary work we conducted on larval performance showed that Hayden’s ringlet larvae can survive on *Poa pratensis* through at least the 4th instar, at which point our larvae entered—and did not survive—diapause. Kentucky bluegrass, *Poa pratensis*, is one of the most popular and widespread turf grass species in the United States (Huff et al., 2003). It is ubiquitous along roadsides and in lawns, occurs in all 50 states, and is highly invasive across the northern Great Plains (DeKeyser et al., 2015). If Kentucky bluegrass is in fact a viable host for the Hayden’s ringlet, it would strongly suggest that host use is not driving endemism in this species. Other environmental factors not considered in this study, including site elevation, temperature, rainfall, tree cover, or other factors may play a far greater role in explaining endemism in the Hayden’s ringlet. The strong impact of geography on genetic differentiation we saw in both our IBD and EEMS results would seem to support this view. More exploration of the niche space inhabited by the Hayden’s ringlet is necessary to more fully rule understand the causes of endemism in this species.

## Conclusions

Despite their restricted range, we found that the Hayden’s ringlet harbors genetic diversity levels comparable to non-endemic species with similar habits. We found strong evidence that the Hayden’s ringlet prefers grasses (*Poa*) over sedges (*Carex*) as a larval host, but work to determine the degree of host specificity in this species remains to be done. Geography, specifically isolation by distance and physical barriers to migration (i.e. mountain ranges and/or regions of poor habitat), appear to be the driving factors producing patterns of population structure in the Hayden’s ringlet. We found no evidence of host-associated divergence, and it does not appear that host specialization is driving either population divergence or range restriction in this endemic species. Instead, population structure in this species has developed largely via genetic drift. Questions remain as to how evolutionary processes unfold in the face of endemism, but in some cases at least, it appears evolutionary dynamics under restricted vs. widespread ranges may not differ as greatly as one might expect.

## Acknowledgments

Thank you to Megan Brady for invaluable assistance with both field collections and preference experiments, to Angelica Traslavina for assistance with DNA extractions, and to both Camden Treat and Daniel Johnson for their extensive assistance cataloguing larval herbivory results. Support and resources from the Center for High Performance Computing at the University of Utah are gratefully acknowledged. This work was funded by the National Science Foundation (NSF GRFP awarded to Amy Springer, fellow 2017239847) and Utah State University.

