## Supplemental Materials for "Considerable genetic diversity and structure despite endemism and limited ecological specialization in the Hayden’s ringlet, *Coenonympha haydenii*"

### Supplement

#### Additional sampling locations

Table S1: Locations of sites that were assessed for the presence of Hayden's ringlets but did not harbor population sizes large enough to sample. Shown are the latitude and longitude of each site, as well as the number of Hayden's ringlets that were observed at each site.

| | Latitude | Longitude | N butterflies observed |
|---|---|---|---|
| AVP | 44.4860 | 110.1307 | 1 |
| GLR | 44.1023 | -110.7483 | 0 |

#### Analyzing leaf images for herbivory with imageJ

We used `ImageJ` version 1.52A (Schneider et al., 2012) to analyze leaf photographs from our larval herbivory trials. Specifically, we analyzed images taken before and after the herbivory trials to determine the total surface area of each leaf consumed during the herbivory trial.

To calculate leaf surface areas, we first set the scale in each image using the `straight` tool in for drawing straight lines. A line was drawn across exactly two blue horizontal grid marks of the graph paper in each image, which corresponds to the known distance of 1.27 cm. To set this known distance as the scale, we used the commands `analyze`→`set scale`, set the parameter `known distance` to 1.27, and the parameter `unit of length` to ”cm”. We then used the `polygon` tool to draw a polygon around only the leaves and grid paper within the image, excluding any writing or any parts of the image beyond the grid paper on which the leaves were placed. The region within the polygon was then duplicated using the the commands `image`→`duplicate`.

Color and contrast were adjusted within the duplicated image using the commands `image`→`type`→`8-bit` to transform the image to grayscale, followed by `image`→`adjust`→`threshold` to transform the image to black and white such that the leaf surfaces were shaded entirely in black and the background was shaded entirely with white. The threshold (bottom sliding bar) was adjusted manually to ensure that the edges of the leaves were precisely highlighted, but no other parts of the image were.

Finally, the area of each leaf was analyzed using the commands `analyze`$\rightarrow$`analyze particles`. We set the parameter `particle size` to a range of 0.01-Infinity to remove noise from the analysis (i.e. particles too small to be leaves were removed). We selected `outlines` from the `show` dropdown menu, which produces an image with the outlines of each particle analyzed in order to double-check that the particles being captured by the analysis corresponded to the shape and location of the leaves in each image. The boxes `exclude on edges`, `display results`, `record starts`, and `include holes` were checked. These parameters serve to exclude any particles touching the edge of the image (as all leaves in our images were centered so particles along edges caould not be leaves), record the starting position (x-y location) of each leaf so it can be located and verified on the original image, and ensure that any areas within leaves that were not shaded black during the `threshold` transformation were still included in area calculations. The result of this analysis is a list of three area values in $cm^2$ corresponding to the surface areas of the three leaves in each image we analyzed. Leaves in each image were always placed in the same order from left to right as follows: *Carex hoodii*, *Campanula rotundifolia*, and *Poa pratensis.*

All leaf images were also manually assessed for signs of herbivory. Specifically, because Hayden's ringlets feed from the margins of leaves rather than skeletonizing tissue from the center, we assessed the margins of each leaf before and after herbivory for jagged edges. Leaves used in the herbivory assays were cut into 1-cm long rectangles prior to herbivory assays so images could be easily coded as either (1) herbivory observed (leaf margins jagged and rectangle sides no longer straight lines) or (0) no herbivory observed (leaf margins intact and no jagged marks present). This manual coding allowed us to isolate changes in leaf surface area due to moisture loss in the leaf tissue over time (i.e. shrinkage) vs. larval herbivory.

#### Construction of the reference contig set with `CD-hit`

Because no reference genome has been constructed for the Hayden's ringlet to date, we constructed a de novo set of reference contigs to align our GBS reads to using the program `CD-hit` version 4.8.1 (Li and Godzik, 2006).

We started with the full set of 347,375,794 demultiplexed reads with poly-G tails removed that remained after filtering to remove PhiX reads. For processing efficiency, we first sorted these reads by individual. This resulted in a single file for each individual containing all the reads that came from that particular individual. We then used `CD-hit` to run a clustering step at 100% match for each individual's set of reads. The result of this step was 287 files containing a list of all unique reads belonging to each individual butterfly. The purpose of this step was to remove reads that were perfect duplicates of one another, thereby reducing file size and processing time downstream.

If we concatenated the files of unique reads from each individual produced above to produce a single file for clustering with `CD-hit` as-is, we would run the risk that `CD-hit` would use solely sequences from the first alphabetically-ordered individual as seed sequences to align other reads to, introducing bias. Instead, we first split the files of unique reads from each individual into approximately 20 files each (70,000 lines or 17,500 reads each). We then concatenated all of the first files split from each individual, followed by all of the second files split from each individual, etc. all the way through the tenth files split from each individual. This ensured that reads from all individual butterflies should be represented in seed sequences during the next clustering step.

To test the sensitivity of percentage match on clustering for our data set, we used a concatenated file containing just the first files split from each individual (approximately 5% of the full concatenated data set). We tested clustering at 80, 90, and 95% match levels. This resulted in 226,668, 349,754, and 419,676 clusters, respectively. We then repeated this sensitivity test using a concatenated file containing the first four files split from each individual (approximately 25% of the full concatenated data set) for 88, 90, and 92% match levels. This resulted in 797,035, 901,765, and 1,006,900 clusters, respectively. We chose to use clustering at 90% match on the full concatenated data set (the first 10 files split from each individual) for our actual clustering to be used in downstream analysis. Any reads that did not cluster at 90% match were then removed from the data set.

Finally, we completed one additional clustering step using only those reads that clustered at 90% match. We clustered these reads again at 80% match and removed all reads that clustered at this level. Reads from the 90% match cluster set that clustered with one another at an 80% match rate or greater were removed because these clusters may represent gene families or duplicated genes. The result of this was a set of 614,359 reference contigs used for alignments in downstream analysis.

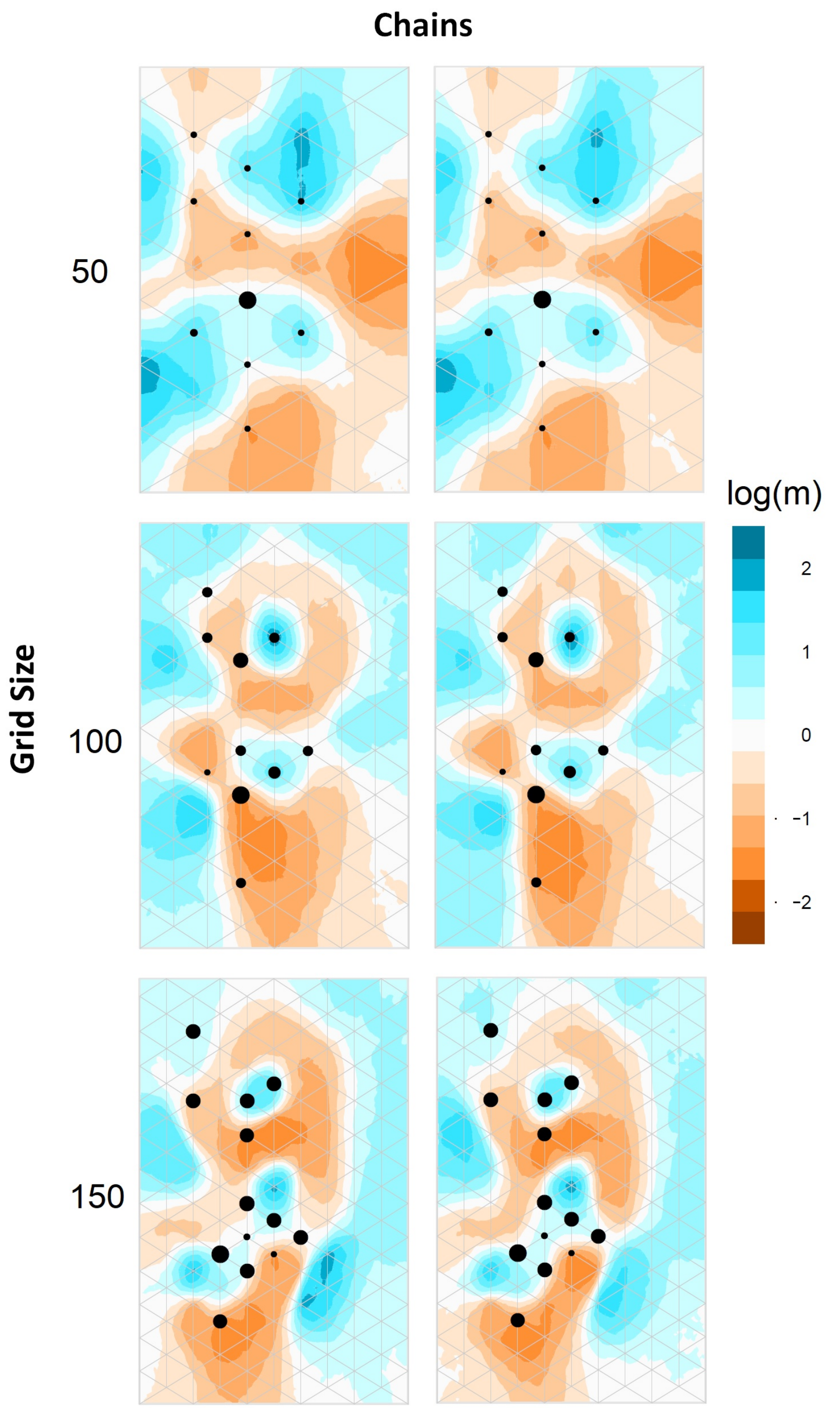


Figure S1: Effect of grid size on `EEMS` results. Two chains each are shown for grid sizes of 50, 100, and 150. Results were highly consistent between chains, and showed an area of reduced relative migration rates between northern and southern populations of Hayden's ringlets across all grid sizes tested.
